# Blood *DDIT4* and *TRIM13* transcript levels mark the early stages of Machado-Joseph disease

**DOI:** 10.1101/2023.10.01.560372

**Authors:** Ana F. Ferreira, Mafalda Raposo, Emily D. Shaw, João Vasconcelos, Teresa Kay, Conceição Bettencourt, Maria Luiza Saraiva-Pereira, Laura Bannach Jardim, Maria do Carmo Costa, Manuela Lima

## Abstract

Machado-Joseph disease (MJD) is a rare late-onset polyglutamine neurodegenerative disease caused by the expansion of a CAG repeat in the *ATXN3* gene encoding the ataxin- 3 (ATXN3) protein. Several studies have identified changes in the abundance of select transcripts and proteins in blood samples of MJD mutation carriers. Here, we aimed to: 1) identify blood transcriptional changes that could be potential biomarkers of MJD from preclinical to symptomatic disease stages; 2) correlate levels of differentially expressed transcripts in blood of MJD carriers with demographic, genetics, and clinical features; and 3) evaluate whether the identified differential abundance of selected transcripts in blood of MJD subjects is preserved in post-mortem brains of MJD patients. Using real-time quantitative PCR, we observed consistent dysregulation of *DDIT4*, *TRIM13* and *P2RY13* transcript levels in blood samples of MJD subjects from different disease stages (from preclinical to symptomatic), and of patients from two cohorts with different backgrounds (Azores and Brazil). Importantly, combined blood *DDIT4* and *TRIM13* transcript levels display a very high accuracy to discriminate MJD carriers in preclinical stage, early-stage patients, and patients with more than five years of disease duration from respective controls (AUC=1.00, AUC=0.96, and AUC=0.90, respectively). Moreover, in the combined group of Azorean and Brazilian patients, blood levels of *P2RY13* transcript correlate with age at onset, and abundance of *DDIT4* and *TRIM13* transcripts correlate with the CAG repeat size of the expanded *ATXN3* allele. In the subgroup of early-stage Azorean patients, blood levels of *TRIM13* transcripts correlate with age at disease onset. Interestingly, abundance of DDIT4, TRIM13 and P2RY13 proteins is also altered in brains of MJD patients. In summary, this work shows that blood *DDIT4* and *TRIM13* transcript levels are potentially blood-based biomarkers of MJD of special usefulness in marking preclinical and early stages of the disease, and points for common dysregulated processes involving *DDIT4*, *TRIM13* and *P2RY13* in the blood and the brain of MJD subjects.

## Introduction

Machado-Joseph disease, also known as Spinocerebellar Ataxia Type 3 (MJD/SCA3; OMIM #109150), is a slowly progressive autosomal dominant ataxia of late onset^1^ predominantly affecting the cerebellar, pyramidal, extrapyramidal, motor neuron and oculomotor systems.^2,3^ The MJD-causative *ATXN3* gene contains a polyQ-encoding CAG tract which, in non-mutated chromosomes ranges from 12 to 44 repeats, consensually surpassing 60 repeats in mutated chromosomes^4^. The overt disease is preceded by a preclinical phase in which MJD mutation carriers already show brain structural changes^5,6^ and may manifest clinical signs such as nystagmus.^7,8^ *ATXN3* encodes ataxin-3 (ATXN3), a deubiquitinating enzyme that is ubiquitously expressed in various cell types of the central nervous system and of peripheral tissues.^9^ Evidence shows that mutated ATXN3 triggers a cascade of pathogenic events leading to several cellular alterations including transcriptional dysregulation.^9,10^ In fact, global changes in gene expression have been identified in peripheral blood samples from MJD carriers,^11–15^ including preclinical subjects.^13,14^ Therefore, tracking significant transcriptional alterations in blood samples of MJD subjects should: i) provide new insights about the peripheral molecular dysfunction in MJD and its association with the primary dysfunction occurring in the brain, and ii) allow the identification of novel minimally invasive molecular biomarkers of MJD, particularly in preclinical stages where clinical scales are devoid of utility. Indeed, several studies have identified changes in levels of transcripts and proteins in peripheral blood samples of MJD carriers, such as alterations in inflammation markers (increase of the pro-inflammatory factor eotaxin),^16^ mitochondrial dysfunction and oxidative stress (decrease of antioxidant capability and excessive production of reactive oxygen species),^17^ accumulation of a common mitochondrial DNA deletion (m.8482_13460del4977) and reduced *BCL2* transcript levels^15^ as well as reduced *BCL2*/*BAX* transcript ratio.^18^

Because clinical trials enrolling MJD subjects at preclinical and early disease stages are expected to demonstrate the maximal efficacy of disease-modifying therapeutic agents^19^, biomarkers of these early disease stages are expected to be the most informative to be assessed over the course of clinical trials.^20^ Increased blood levels of the general biomarker of neurodegeneration - the neurofilament light chain (NfL) - and of mutant ATXN3 are currently the most promising blood biomarkers for MJD, altought further studies to establish their value as progression, proximity-to-onset, and putative treatment-response markers are still needed, justifying the need to explore additional complementary biomarkers.

Hence, in this study, we sought to identify transcriptional changes in peripheral blood of MJD subjects that could be potential molecular biomarkers of MJD from preclinical to symptomatic phases and, ideally, of disease progression. To investigate the behavior of such blood-based transcriptional changes in the primarily affected tissue in MJD, we further assessed the expression pattern of the most promising candidates at the transcript and protein levels in post-mortem brain samples of MJD patients and controls.

## Material and methods

### Blood samples and clinical data

A total of 83 blood samples from MJD patients and 19 from preclinical MJD subjects were used in this study. The present study included 59 patients representing several stages of the disease, 39 from the Azorean cohort and 20 from the Brazilian cohort. A subgroup of 36 patients from the Azorean cohort in the early symptomatic stage of the disease (EP), defined as having five or less years of disease duration, were also studied; this group is formed by 12 patients from the group of Azorean patients mentioned above (n=39), plus 24 additional independent Azorean patients. The five year cut-off used to define the early patients was based on the natural history data for MJD showing that the mean score of disease progression (defined by gait impairment) progresses very slowly during the first five years.^21^ All patients underwent a neurological evaluation by experienced neurologists in MJD (LJ and JV), and for a subset of patients the score of the Neurological Examination Score for Spinocerebellar Ataxia (NESSCA) was assessed.^22^ Age at onset was defined as the age of appearance of the first symptoms (gait ataxia and/or diplopia) reported by the patient or a close relative. Disease duration was defined as the time elapsed between age at onset and age at neurological evaluation during which blood was collected. Nineteen Azorean preclinical subjects (PC) were also enrolled in the study after being confirmed as carriers of the *ATXN3* mutation, in the context of the Azorean Genetic Counseling and Predictive Test Program offered by the Regional Health System (Azores, Portugal). For preclinical subjects, the number of years between the age at neurological evaluation/blood collection and the predicted age at disease onset (years to onset, time) were calculated as previously described by Raposo and colleagues.^23^ The determination of the number of CAG repeats at the *ATXN3* locus was performed for MJD subjects as described by Bettencourt and colleagues.^24^ Additionally, blood samples from 98 apparently healthy controls from the same two populational origins (78 from Azores and 20 from Brazil) as the MJD subjects were also included in this study. In addition, the controls were molecularly excluded for the *ATXN3* mutation using the above-mentioned protocol.^24^ Characterization of the MJD subjects and controls used in this study are shown in Supplementary Table 1. This study was approved by local Ethics Committees from the participating institutions. All participants provided written informed consent.

### Post-mortem human brain samples

To investigate the expression patterns of a subset of relevant genes, post-mortem flash frozen brain samples from deidentified molecularly confirmed MJD patients (n=5) and controls (n=9) obtained from the University of Michigan Brain Bank were used. Samples from two brain regions severely affected in MJD (dentate cerebellar nucleus (DCN) and pons)^25^, and from a region minimally affected by the disease (frontal cortex)^26,27^ were used in this study. The CAG repeat number of all post-mortem brain samples was determined at Laragen Inc. (Culver City, CA) by DNA fragment analysis as described by Ashraf and collegues^28^. Demographic, genetic, and clinical information of the MJD patients and controls from whom brain samples were used in this study are shown in Supplementary Tables 1-2. The use of human post-mortem brain samples in this study was approved by the University of Michigan Biosafety Committee (IBCA00000265) and by the Ethics Committees of the University of the Azores (Parecer 26/2018).

### Experimental design

An overview of the experimental workflow used in this study is displayed in Fig. 1 and the characterization of the MJD subjects and controls used in each step is shown in Supplementary Table 1. Briefly, 28 peripheral blood RNA samples from the Azorean cohort (group #1) were used to perform a gene expression microarray study (Fig. 1A). To identify dysregulated genes in MJD subjects comparing with healthy controls and to detect differences between the preclinical and the symptomatic stages of the disease, gene expression levels were compared pairwise between the three groups (preclinical subjects vs. controls, patients vs. controls, and preclinical subjects vs. patients). Among the dysregulated genes in blood (Benjamini-Hochberg false discovery rate adjusted *p*-value (*q*-value) lower than 0.05), 24 known to be expressed in the human brain (Expression Atlas database^29^) and without copy number variation (Database of Genomic Variants^30^) were selected for validation by quantitative real-time PCR (qPCR) in the same samples used in the microarray experiment (group #1, Fig. 1B). After technical validation, a set of genes that maintained the same direction of effect (in at least two comparisons) in relation to the microarray was selected if displaying significant differences (*p*<0.05) (n=8), or based on previous data showing that these genes belong to a gene family dysregulated in polyQ diseases (n=4).^31–37^ The expression levels of the selected genes (n=12) were then assessed in independent Azorean blood samples from group #2 (Fig. 1C.1). Next, the expression of five of these 12 genes (showing differential expression in patients compared to controls from group #2) was evaluated in patients and controls from a Brazilian cohort (group #3) (Fig. 1C.2). *P2RY13* transcript levels (purinergic receptor P2Y13, TaqMan assay ID Hs03043902_s1), prior identified by our group as a potential biomarker of MJD,^11^ was also analyzed in the Brazilian cohort, group #3 (Fig. 1C.2). Expression levels of the three genes that showed a consistent dysregulation pattern in blood samples of Azorean and Brazilian patients compared to controls were further evaluated in: i) RNA from blood of MJD Azorean subjects at early stages of disease (preclinical subjects and early patients) and age- and sex-matched paired controls (group #4) (Fig. 1D); and ii) RNA and protein lysates from post-mortem brain tissue from MJD patients and controls (Fig. 1E).

**Figure 1.**
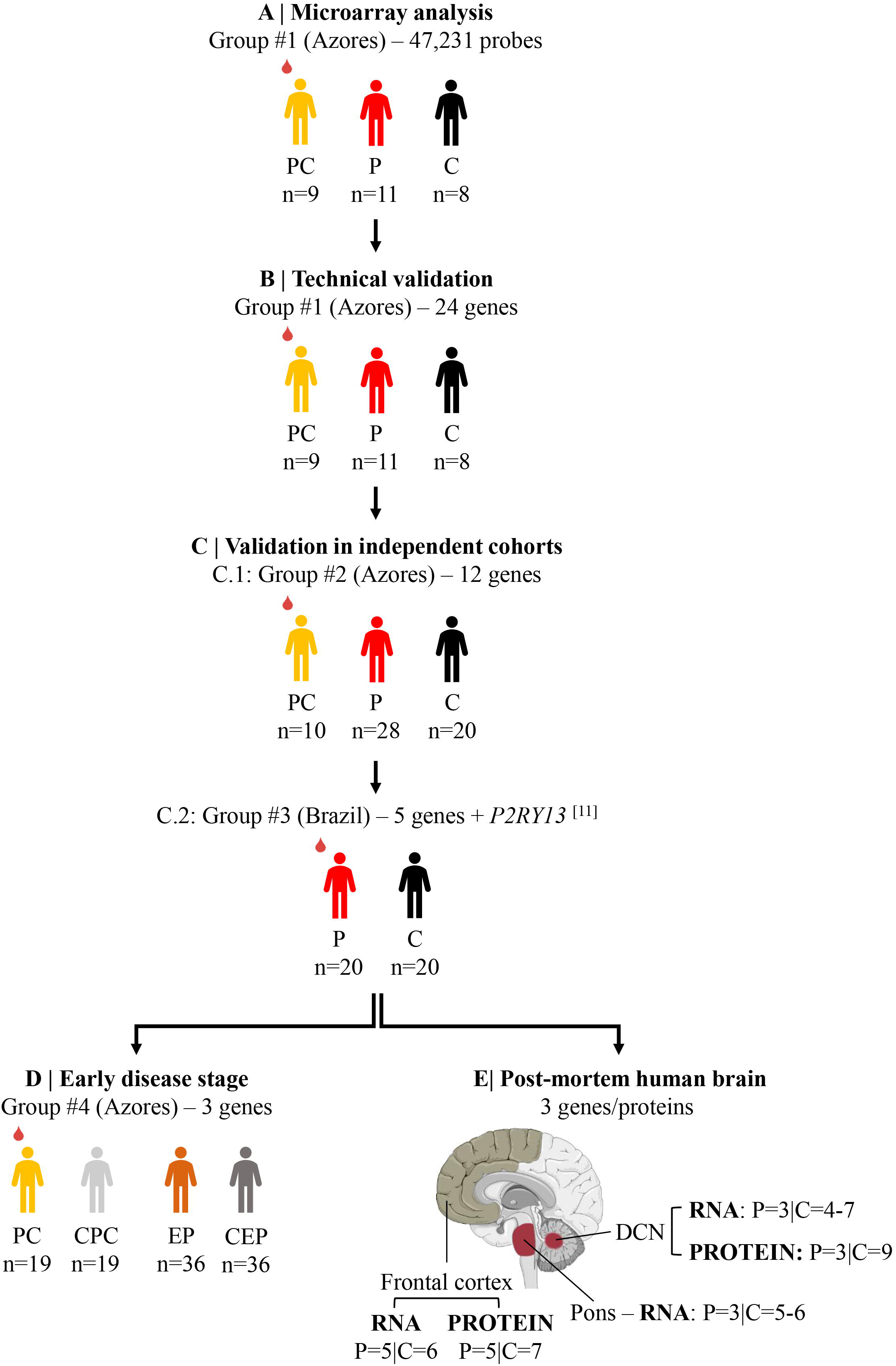
Overview of the experimental workflow used in this study. Peripheral blood samples from MJD subjects and controls from the Azorean cohort were used in **(A)** a genome-wide expression microarray analysis and **(B)** technical validation by qPCR. **(C)** The expression levels were evaluated in two independent cohorts: **(C.1)** The expression levels of a set of dysregulated genes were assessed in a set of Azorean samples; and **(C.2)** genes considered as the most promising disease markers were evaluated in a set of patients and controls of the Brazilian cohort. **(D)** The expression levels of the validated genes were assessed in a set of Azorean PC subjects and early patients. **(E)** The expression of the validated genes was evaluated at the transcript and protein levels in post-mortem brain samples from MJD patients and controls. Human brain figure was partly generated using Servier Medical Art, provided by Servier, licensed under a Creative Commons Attribution 3.0 unported license). PC=preclinical subjects (yellow); EP=early patients (orange); P=patients (red); C=controls (black); CPC=controls of PC subjects (light grey); CEP=controls of EP (dark grey).

### RNA extraction and cDNA synthesis

Total RNA extracted from peripheral blood samples of MJD carriers and heathy controls, as described by Raposo *et al.*^11^, was available for this study. Total blood cDNA was generated from 0.5µg of total RNA using High-Capacity cDNA Reverse Transcription Kit (Applied Biosystems). Total RNA extracted from three brain regions (pons, DCN and frontal cortex) from post-mortem brain samples of MJD patients and controls was extracted using Trizol (Invitrogen) and purified using the RNeasy mini kit (Qiagen) following the manufacturer’s instructions. Brain RNA concentration and integrity were assessed using the Agilent 2100 Bioanalyzer (Supplementary Table 2). Total brain cDNA was generated from 1.25µg of total RNA using the iScript cDNA synthesis kit (BIORAD).

### Microarray analysis

Microarray analysis was conducted to identify transcriptional alterations in peripheral blood samples across the three biological comparison groups: preclinical subjects vs. controls, patients vs. controls, preclinical subjects vs. patients (Fig. 1A; Supplementary Table 1). RNA was amplified, biotin-labeled and directly hybridized to the Illumina HumanHT-12 v4.0 Expression BeadChip (47,231 probes) (Illumina, San Diego, CA, USA). Slides were scanned using the Illumina iScan and the signal was extracted and normalized using the GenomeStudio software. All array procedures were conducted at the UCL Genomics facility (London, UK). Microarray probes related to mRNAs encoding unmapped genes, non-coding genes, pseudogenes and open-reading frames were excluded from further analysis. Mayday 2.9 software^38^ was used to compare the gene expression levels between the three comparison groups (*q*<0.05). The microarray data for the 24 genes selected for qPCR validations are shown in the Supplementary Table 3.

### Quantitative real-time PCR

qPCR experiments (Fig. 1B-E; Supplementary Table 1) were conducted using the TaqMan Gene Expression Master Mix and prevalidated TaqMan Gene Expression Assays (Applied Biosystems, TaqMan assays IDs are shown in Supplementary Table 4), on a StepOnePlus Real-Time PCR System (Applied Biosystems). Each sample was run in triplicate alongside with the reference gene. In the experiments for validations of expression levels from peripheral blood samples from groups #1-3 (Fig. 1B-C), *PPIB* (Hs00168719_m1) was used as reference gene.^11,39^ In the qPCR experiments using blood RNA-derived samples from group #4 (Fig. 1D) and from post-mortem human brain samples (Fig. 1E), *TRAP1* (Hs00972326_m1)^39^ and *GAPDH* (glyceraldehyde-3- phosphate dehydrogenase, Hs02758991_g1) were used as reference genes, respectively. Relative expression values were normalized to the respective reference gene using the 2^-ΔCt^ method^40^ in DataAssist v3.0 software (Applied Biosystems).

### Western Blotting

Protein extracts from DCN and frontal cortex tissues were obtained by homogenization of brain tissue in cold phosphate-buffered saline (PBS) containing protease inhibitor cocktail (cOmplete Protease Inhibitor Cocktail, Roche) and phosphatase inhibitors (PhosSTOP, Roche), followed by sonication and centrifugation. The supernatant was collected (PBS-soluble fraction) and stored at -80°C. The pellet was resuspended in 1% sodium lauroyl sarcosinate (sarkosyl, Sark)/PBS, sonicated, centrifuged and the supernatant (sarkosyl-soluble fraction) was stored at -80°C. Total protein concentrations from PBS-soluble (soluble) and sarkosyl-soluble (insoluble) fractions were both assessed using the BCA method (Pierce BCA Protein Assay Kit; Thermo Fisher Scientific). Total protein lysates (50μg) were resolved on 4-20% SDS-PAGE gel, and corresponding polyvinylidene difluoride membranes were incubated overnight at 4°C with various antibodies: rabbit anti-DDIT4 (1:1000, ab191871, Abcam), rabbit anti-TRIM13 (1:1000, MBS7045610, MyBioSource), rabbit anti-P2RY13 (1:1000, MBS7002276, MyBioSource), rabbit anti-MJD antibody (1:10000^41^), and mouse anti-GAPDH (1:30000, MAB374, Millipore). Bound primary antibodies were visualized by incubation with peroxidase-conjugated goat anti-rabbit or anti-mouse antibodies (1:10000, Jackson Immuno Research Laboratories), followed by reaction with ECL-plus reagent (Western Lighting, PerkinElmer) and exposure to autoradiography films (Fig. 1E). Film band intensity was quantified by densitometry using ImageJ.

### Statistical analysis

Descriptive statistical analysis was used to explore the demographic, genetic and clinical data of MJD subjects. Normal distribution of continuous variables was tested using the Shapiro-Wilk test, and nonparametric tests were used when normality assumption was violated. The comparison between two categorical variables (sex/groups) was tested using a chi-square test, and the comparison of age at blood collection and the expanded CAG repeat tract between groups was tested using a Kruskal-Wallis test or a Mann-Whitney U test (two-tailed).

#### Comparisons of expression levels between groups

*Blood samples* – the comparison of transcripts levels between groups in Azorean cohorts (technical validation, group #1 and validation in independent cohorts, group #2) was performed using a Generalized Linear Model with Bonferroni correction for multiple comparisons whenever applicable (covariates: age at blood collection and/or the size of the expanded CAG repeat) or using a Mann-Whitney U test (two-tailed) in Brazilian cohort (group #3). The comparison of transcript levels between preclinical subjects or early patients and age- and sex-matched paired controls (Azorean cohort, group #4) was performed using a Wilcoxon Signed-rank test (two-tailed). Additionally, a receiver operating characteristic (ROC) curve analysis was performed to determine the ability of each transcript abundance to differentiate: i) preclinical subjects and early patients from age- and sex-matched paired controls (group #4); and ii) patients with more than five years of disease duration from controls (groups #1-3). In the case of the *P2RY13* gene, the latter analysis (patients >5 years of disease duration) was conducted using the gene expression data from Azorean MJD patients and controls obtained from a previous study^11^ and data generated in the present study for group #3. ROC curve analysis was also performed for the combination of the genes using the predicted probabilities obtained by multiple logistic regression. ROC curve accuracy was measured by the area under the curve (AUC), 95% confidence interval (CI), *p-*value; AUC value varies between 0 to 1: 0.5 is equivalent to randomly classifying subjects and 1.00 is equivalent to perfectly classifying subjects. Youden Index^42^ was used to determine the optimal cut-off value for each transcript abundance to differentiate the three groups of MJD subjects (preclinical, early patients and patients > 5 years of disease duration) from their respective controls. *Post-mortem human brain samples* – the comparison of transcript and protein levels between groups was evaluated using a Generalized Linear Model, adjusted for the age at death when this variable had an effect in the model.

#### Association with demographic, genetic and clinical MJD subjects’ features

*Blood samples* – the relationship between transcript levels and MJD subjects’ features was evaluated using a Spearmańs rank correlation (two-tailed); a partial correlation (two-tailed) was used to adjust for covariates (expanded CAG repeat and/or disease duration), when necessary. The relationship between transcript levels and MJD subjects features was tested in two different patients’ groups: i) patients representing several stages of the disease (groups #1-3); and ii) early patients (group #4). In the case of the *P2RY13* gene, the first analysis was conducted using the gene expression data from Azorean patients obtained from a previous study^11^ and data generated for group #3 in the present study.

GraphPad Prism version 8.0.0 for Windows (GraphPad Software, San Diego California USA) was used to identify the extreme outliers (ROUT method Q=1%) that were then excluded from data analysis, to perform the ROC curve analysis, and to generate all the graphs. Statistical procedures were performed using IBM SPSS Statistics for Windows, version 25 (IBM Corp., Armonk, N.Y., USA). Statistical significance was set at *p*<0.05.

### Data availability

Most data are available in this manuscript and in the supplementary material. The raw data of the genome-wide microarray that support the findings of this study are available from the corresponding authors, upon reasonable request.

## Results

### *DDIT4*, *TRIM13* and *P2RY13* transcript levels are altered in blood samples from two cohorts of MJD patients

To identify transcriptional changes in MJD, a gene expression microarray analysis was performed using peripheral blood RNA samples from preclinical MJD subjects, MJD patients, and controls (group #1, Fig. 1). About 9400 probes (of a total of 47,231 probes probes) corresponding to 5209 genes passed the quality control checks. Comparison of blood transcript levels between the three biological groups (preclinical subjects vs. controls, patients vs. controls, preclinical subjects vs. patients) identified 77 genes with significant differences (*q*<0.05) in at least one comparison. Among these dysregulated genes, 24 genes were selected to be technically validated by qPCR in the same blood RNA samples used in the microarray (group #1, Fig. 1) (Supplementary Table 3). Of these 24 genes, *DDIT4* (DNA Damage Inducible Transcript 4), *TRIM13* (tripartite motif containing 13), *S100P* (S100 calcium binding protein P), *SVBP* (small vasohibin binding protein) and *UPS49* (ubiquitin specific peptidase 49) maintained their dysregulation pattern in independent samples from Azorean MJD patients (group #2, Fig. 1) and were then further analyzed in samples from MJD patients and controls from an independent Brazilian cohort (group #3, Fig. 1) (Fig. 2; Supplementary Table 4). Levels of blood *P2RY13* transcripts, which were previously shown to be increased in Azorean MJD patients,^11^ were also investigated in samples from the Brazilian cohort (Fig. 2). Because *P2RY13* failed to pass the quality control of the microarray experiment, no expression data from this experiment was determined. In summary, the three genes (*DDIT4*, *TRIM13*, and *P2RY13*) displayed a consistent expression behavior in blood samples from MJD patients compared to controls in the two independent cohorts of different populational origins (Azores and Brazil) (Fig. 2): the levels of *DDIT4* were decreased, whereas the levels of *TRIM13* and *P2RY13* were increased in MJD patients. Moreover, *TRIM13* transcript levels were increased in Azorean preclinical MJD subjects comparing with the respective controls (Fig. 2).

**Figure 2.**
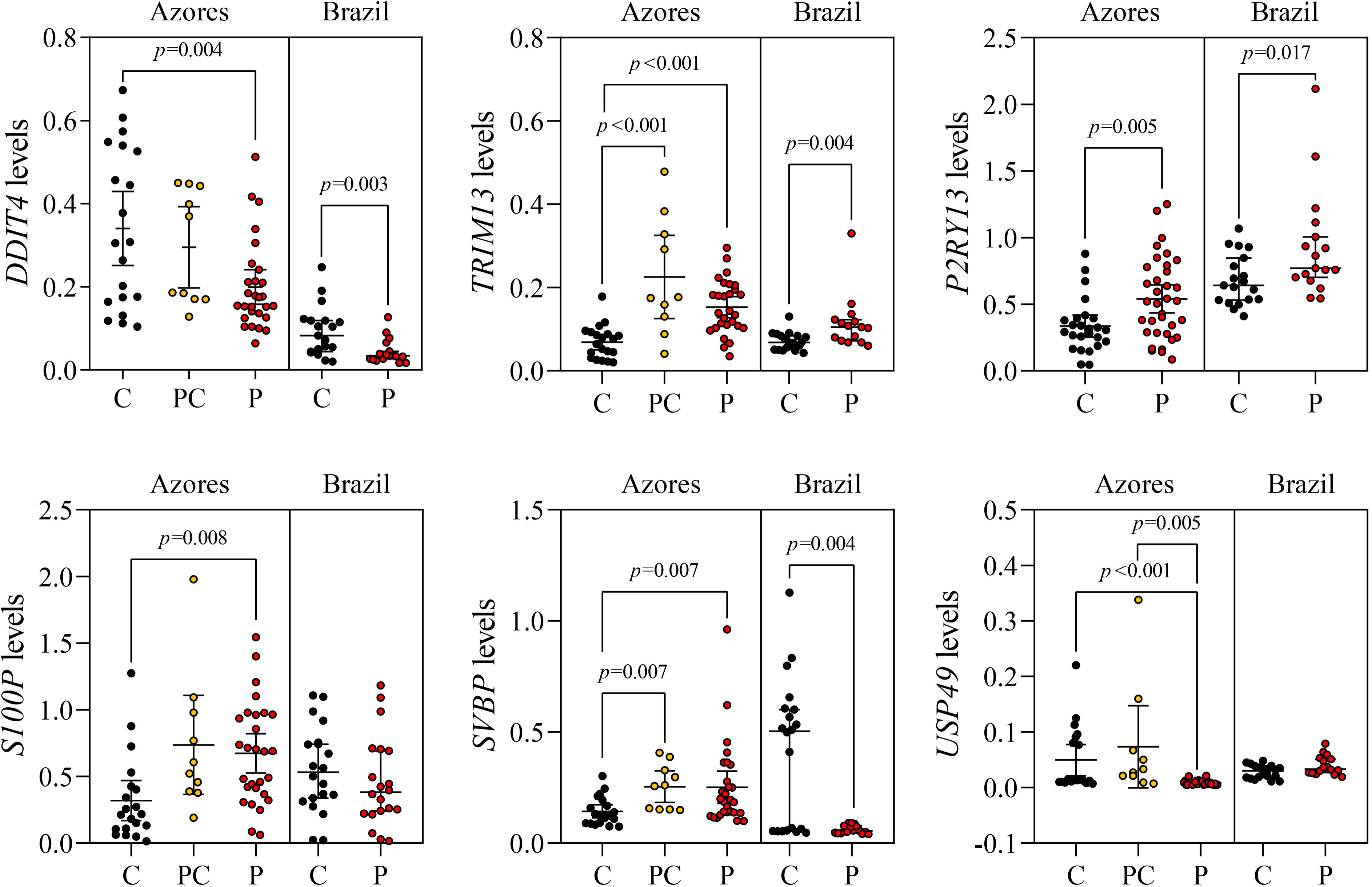
Blood transcript levels of *DDIT4*, *TRIM13* and *P2RY13* are consistently dysregulated in two independent MJD cohorts. Gene expression levels of *DDIT4*, *TRIM13, P2RY13 S100P*, *SVBP* and *USP49* in peripheral blood samples from MJD carriers (PC, preclinical subjects (yellow dots), and P, patients (red dots) and controls (C, black dots) from Azorean cohort (group #2) and MJD patients (P, red dots) and controls (C, black dots) from Brazilian cohort (group #3). Dots represent transcript levels for each individual. Graphs show the mean (Azores) or median (Brazil) and the 95% confidence interval of the 2^-ΔCt^ values.

### Blood transcript levels of *DDIT4* and *TRIM13* correlate with the CAG repeat size of patients and *P2RY13* levels with the age at onset

Blood transcript levels of *DDIT4*, *TRIM13* and *P2RY13* were correlated with several demographic, genetic, and clinical features of MJD patients (groups #1-3, Fig. 1; Supplementary Table 5). *DDIT4* and *TRIM13* levels inversely correlated with the CAG repeat number of the expanded *ATXN3* allele (rho=-0.290 and *p*=0.027, and rho=-0.330 and *p*=0.015, respectively), and *P2RY13* levels positively correlated with the age at disease onset (rho=0.366 and *p*=0.004) (Supplementary Table 5).

### Blood levels of *DDIT4*, *TRIM13* and *P2RY13* transcripts are altered in the initial stages of MJD

To determine the potential use of *DDIT4*, *TRIM13* and *P2RY13* transcript levels as blood-based biomarkers of initial stages of MJD, their abundance was evaluated in preclinical subjects, early-stage patients (< 5 years of disease duration) and age- and sex-matched paired controls (early disease stage, group #4, Fig.1D) (Fig. 3). *DDIT4* levels were decreased whereas *TRIM13* and *P2RY13* levels were increased in both preclinical subjects and early-stage patients (Fig. 3). These results observed for early-stage patients (group #4, Fig. 1D) are in accordance with those obtained for the cohorts of MJD patients from the Azores and Brazil (groups #2 and 3, respectively, Fig.1C) that comprise subjects in a wider range of disease stages (up to 25 years after onset) (Fig. 2). Furthermore, when correlating the transcript levels of these three genes with the expanded CAG repeat size and the clinical features of early-stage patients, *TRIM13* levels inversely correlated with the age at disease onset (rho=-0.472, *p*=0.004) (Supplementary Table 6), suggesting that the differential blood *TRIM13* expression may be associated with disease severity.

**Figure 3.**
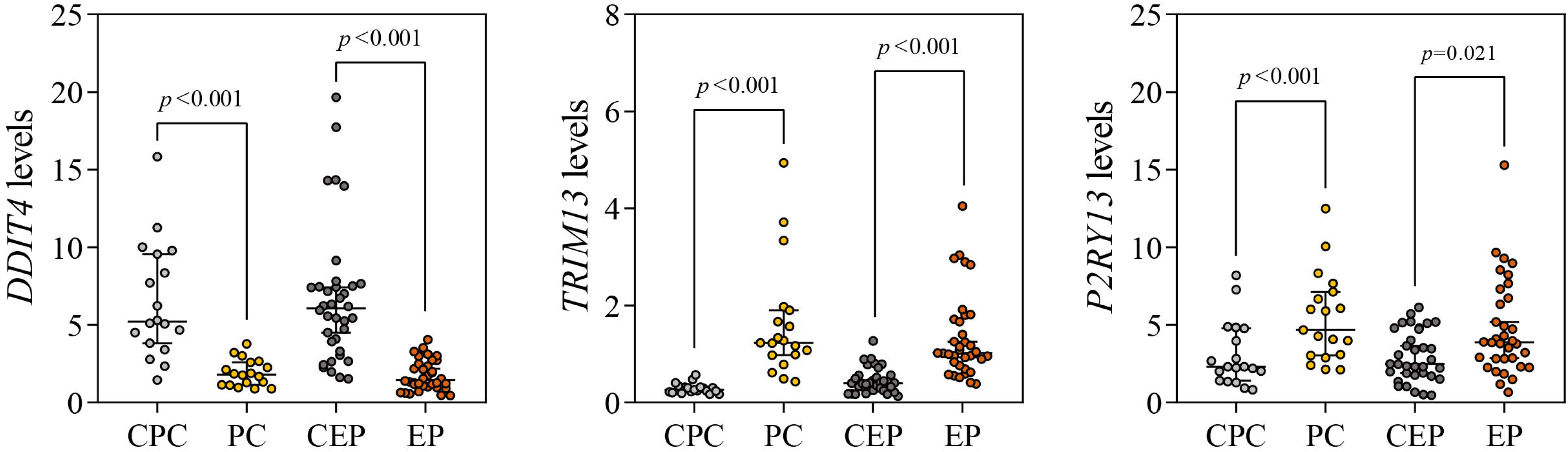
Blood transcript levels of *DDIT4*, *TRIM13* and *P2RY13* are dysregulated in initial stages of the disease. Genes expression levels in peripheral blood samples from preclinical (PC, yellow dots) subjects and early patients (EP, orange dots), and age- and sex-matched paired controls (CPC, controls of preclinical subjects (light grey dots); CEP, controls of EP (dark grey dots)) from group #4. Dots represent transcript levels for each individual. Graphs show the median and the 95% confidence interval of the 2^-ΔCt^ values.

### Blood transcript levels of *DDIT4* and *TRIM13* differentiate MJD subjects from controls with high accuracy

Next, the ability of blood transcript levels of *DDIT4*, *TRIM13* and *P2RY13* to differentiate MJD subjects from matched controls was investigated in various stages of the disease: i) in MJD subjects at initial stages of the disease (preclinical subjects and early-stage patients, group #4), and ii) in MJD patients with more than five years of disease duration (Fig. 4). *TRIM13* and *DDIT4* levels differentiated with very high accuracy preclinical subjects from controls (*TRIM13*: AUC=0.99 (0.97-1.00), *p*<0.001; *DDIT4*: AUC=0.94 (0.86-1.00), *p*<0.001), as well as early-stage patients from controls (*TRIM13*: AUC=0.92 (0.85-0.98), *p*<0.001; *DDIT4*: AUC=0.93 (0.87-0.98), *p*<0.001) (Fig. 4). On the other hand, *P2RY13* levels differentiated preclinical subjects and early-stage patients from controls with moderate (AUC=0.78 (0.63-0.93), *p*=0.004) and low accuracy (AUC=0.68 (0.55-0.81), *p*=0.010), respectively (Fig. 4). Importantly, the combination of *DDIT4* and *TRIM13* levels perfectly differentiated preclinical subjects from controls (AUC=1.00 (1.00-1.00), *p*<0.001). In addition, the combination of *DDIT4* and *TRIM13* levels also improved the ability to differentiate early-stage patients from controls (AUC=0.96 (0.92- 1.00), *p*<0.001) (Fig. 4), while no improvement was seen when *P2RY13* abundance was added to the other two genes (AUC=0.96 (0.91-1.00), *p*<0.001). Noteworthy, blood *TRIM13* transcript levels further differentiated with good accuracy MJD patients with a disease duration longer than five years from controls (AUC=0.85 (0.77-0.93), *p*<0.001), and the combination of *DDIT4* and *TRIM13* levels improved the ability to differentiate these two groups (AUC=0.90 (0.83-0.96), *p*<0.001) (Fig. 4). The optimal cut-off values for *DDIT4*, *TRIM13*, and *P2RY13* transcript levels to differentiate MJD subjects in several disease stages from controls are summarized in Supplementary Table 7.

**Figure 4.**
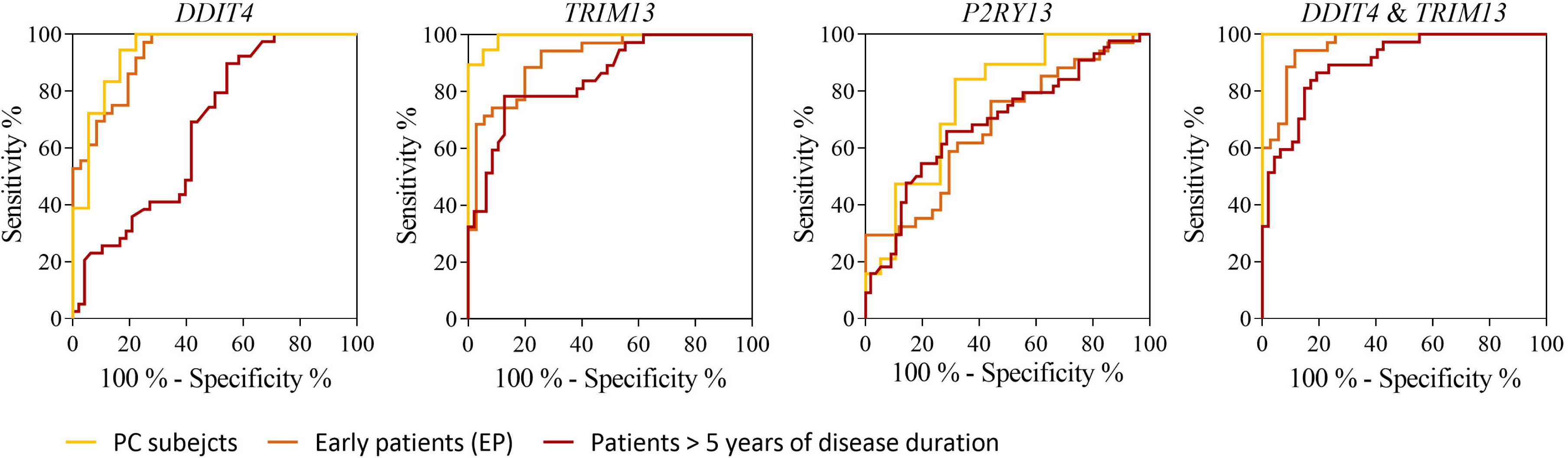
Blood transcript levels of *DDIT4* and *TRIM13* differentiate with very high accuracy MJD subjects from controls. Receiver operating characteristics curves show the ability of blood transcript levels of *DDIT4*, *TRIM13*, *P2RY13* and the combination of *DDIT4* and *TRIM13* levels to differentiated between MJD subjects in different stages of the disease (PC, yellow line; EP, orange line; patients with > 5 years of disease duration, dark red line) and matched controls.

### DDIT4, TRIM13 and P2RY13 protein levels are dysregulated in brain samples from MJD patients

The expression patterns of *DDIT4*, *TRIM13*, and *P2RY13* were assessed at the transcript and protein levels in a set of post-mortem brain samples from MJD patients and controls. The amount of *DDIT4*, *TRIM13* and *P2RY13* transcripts in the three analyzed brain regions (pons and DCN that are severely affected in MJD, and frontal cortex that shows a lower degree of degeneration in MJD, as previously referred) was similar in MJD patients and controls (Fig. 5).

**Figure 5.**
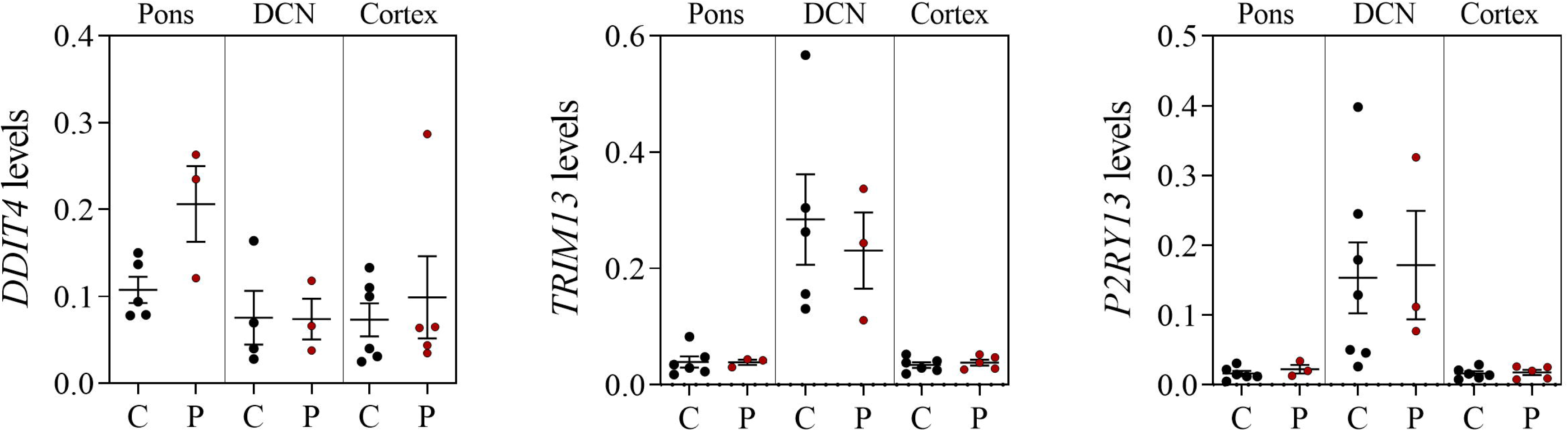
*DDIT4*, *TRIM13* and *P2RY13* transcript levels are similar in post-mortem brain of MJD patients and controls. Transcript levels in pons, dentate cerebellar nucleus (DCN) and frontal cortex (cortex) from MJD patients (P, dark red dots) and controls (C, black dots). Dots represent transcript levels for each individual. Graphs show the mean and the standard error mean of the 2^-ΔCt^ values.

The abundance of DDIT4, TRIM13 and P2RY13 proteins assessed in the DCN and in the frontal cortex showed significant alterations in MJD patients compared with control subjects (Fig. 6; Supplementary Fig. 1): i) soluble DDIT4 levels were increased in the frontal cortex (*p*=0.037); ii) soluble TRIM13 levels were increased in the DCN (*p*=0.026); and iii) soluble P2RY13 levels were decreased in the frontal cortex (*p*=0.010), whereas insoluble protein levels were increased in the DCN (*p*=0.021).

**Figure 6.**
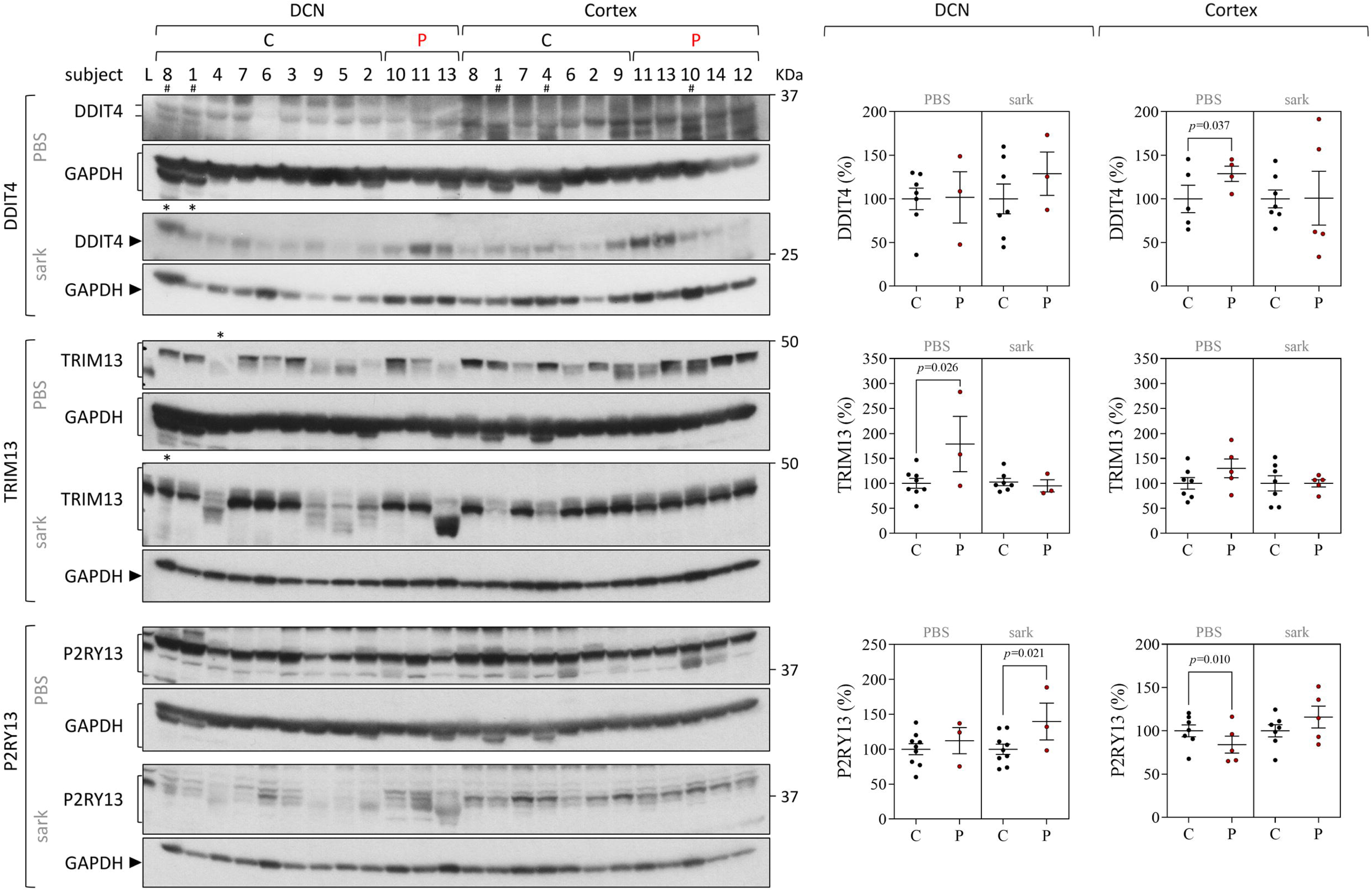
DDIT4, TRIM13 and P2RY13 protein levels are altered in post-mortem brain of MJD patients. PBS-soluble (soluble) and sarkosyl-soluble (insoluble) protein fractions were analyzed in dentate cerebellar nucleus (DCN) and frontal cortex (cortex) from MJD patients (P, dark red dots) and controls (C, black dots). Left panels show immunoblots of indicated proteins in soluble and insoluble protein fractions. Right panels display quantification of band intensity, with values normalized to GAPDH. Dots represent protein levels for each individual. Graphs show the mean and the standard error mean of protein relative to controls. L, protein molecular weight ladder loaded in the protein gels. Samples labeled with a symbol were excluded from statistical analysis due to (*) gel irregularities or (#) quantification difficulties.

## Discussion

This study aimed to identify transcriptional changes in peripheral blood samples from MJD subjects that could be used as prospective biomarkers of MJD, in particular of its initial stages. Data generated by our unbiased genome-wide expression microarray study confirmed the existence of global transcription dysregulation in blood samples from MJD subjects, which had been previously reported (our microarray study identified common dysregulated gene expression with previous studies, such as the Charcot-Leyden crystal galectin (*CLC*) and the Peptidase inhibitor 3(*PI3*) genes).^11,14^ To investigate whether the transcriptional alterations found at the periphery are conserved in the tissue primarily affected in MJD, the transcript and protein levels of three most promising candidates were assessed in post-mortem brain samples from MJD patients and controls.

Building up on the cross-sectional blood-based microarray data of MJD subjects and controls conducted in this study and on a previous gene expression study,^11^ three genes – *DDIT4*, *TRIM13* and *P2RY13* – were identified as being consistently dysregulated in blood samples of MJD carriers in various stages of disease, including preclinical subjects and early-stage patients. Our results showed that individual blood transcript levels of *DDIT4* and *TRIM13* were able to differentiate with very high accuracy (AUC>0.90) both preclinical subjects and early-stage patients from age- and sex-matched paired controls. Remarkably, the combination of *DDIT4* and *TRIM13* blood transcript levels fully differentiated preclinical subjects from controls (AUC=1.00), and further differentiated with high accuracy patients with more than five years of disease duration from controls (AUC=0.90). To date, the levels of NfL^43–47^ and of mutant ATXN3 protein^46,48,49^ have been considered the blood-based molecular biomarkers that would better differentiate MJD subjects from control subjects and correlate with clinical and genetic features of the disease. However, the use of blood NfL and mutant ATXN3 levels as biomarkers of MJD shows some limitations: i) blood NfL levels are not a specific biomarker of MJD but a general biomarker of neurodegeneration, and are not only a reflex of disease processes but also of other factors such as age, body-mass index, diabetes and cardiovascular risk factors;^50^ and ii) blood mutant ATXN3 levels may not be useful to assess the efficacy of disease-modifying therapeutic agents other than the ones targeting *ATXN3* gene products. In addition, only some of the aforementioned NfL studies assessed whether blood-based molecular alterations differentiate premanifest subjects^43–45,47,49^ and patients at early stages of the disease^43^ from controls, limiting the comparisons of our current study with these previous studies. Noteworthy, prior to disease onset, blood *DDIT4* and *TRIM13* transcript levels, individually or combined, differentiated MJD carriers from healthy controls with higher accuracy (*DDIT4*: 94%, *TRIM13*: 99% and *DDIT4+TRIM13*: 100%, respectively) than serum and plasma NfL levels (79-89%).^43,44,47^ Concerning the early disease stage, the combination of *DDIT4* and *TRIM13* transcript levels and serum NfL levels^43^ differentiate early-stage patients from controls with similar accuracy (96% and 98%, respectively). These findings highlight the potential of *DDIT4* and *TRIM13* transcript levels as blood-based biomarkers of MJD, in particular for the preclinical and early symptomatic stages of the disease.

Reinforcing the hypothesis that such differences in expression are meaningful and indeed linked to the disease process, the expression levels of *DDIT4*, *TRIM13* and *P2RY13* genes correlated with patients’ age at onset and/or the number of CAG repeats in the expanded allele supporting a potential role in MJD pathogenesis. While the number of assessed post-mortem brain samples in this study was low, the proteins encoded by these three genes were altered in brains from MJD patients compared to controls (no changes were found at the mRNA levels), further indicating the involvement of these proteins in MJD- associated mechanisms.

*DDIT4* encodes the DNA-damage-inducible transcript 4 protein (mainly localized in the cytoplasm) which is mainly expressed in T-cells and macrophages, and in astrocytes.^51^ DDIT4 protein is induced by cellular stress playing a role in regulation of cell growth, proliferation and survival by inhibiting the activity of mTORC1 (mammalian target of rapamycin complex 1).^52–54^ Prolonged DDIT4 expression is associated with apoptosis, excessive production of reactive oxygen species and inflammation activation ultimately leading to tissue damage.^55^ In this study, blood *DDIT4* transcript levels were decreased in MJD carriers and inversely correlated with the size of the expanded CAG repeat of patients. The results observed in the blood samples of MJD subjects are in line with several studies that show impaired mitochondrial function in blood of MJD,^17,18^ since reduced abundance of DDIT4 protein can result in loss of mitochondrial integrity leading to increased inflammatory response.^56,57^ Interestingly, while DDIT4 protein levels in the affected DCN and pons were similar in patients and controls, the less-affected frontal cortex of patients showed higher soluble DDIT4 protein levels than controls possibly suggesting that cell death is being triggered. The hypothesis that the apoptotic process is activated in the less affected frontal cortex and not in the severely affected DCN and pons of MJD patients is in line with the results recently reported by our group that show increased anti-apoptotic signs (BCL2/BAX ratio) in the DCN and pons from the same MJD patients used in this study, suggesting that the most affected neuronal and glial cells are most probably dead and absent at the end stage of the disease^15^.

To the best of our knowledge, *TRIM13* encoding the Tripartite motif containing 13 protein has not yet been associated with any neurodegenerative disease. *TRIM13* is expressed in immune cells and in neuronal and glial cells (oligodendrocytes, microglial and astrocytes).^51^ TRIM13 protein is a positive regulator of nuclear factor-kappa B (NF-kB) signaling, inducing the production of cytokines and inflammatory mediators.^58^ TRIM13 protein has also been reported to be involved in other processes altered in MJD including the elimination of endoplasmic reticulum (ER)-associated protein degradation substrates,^59^ regulation of autophagy,^60,61^ and cell death during ER stress.^62^ Tripartite motif (TRIM) family members are crucial players in eliminating misfolded proteins,^63^ and some of these proteins have been associated with neurodegenerative diseases, namely TRIM19 in SCA1^34^ and TRIM11 in Parkinson’s and Alzheimer diseases.^36,37^ Comparing with controls, abundance of blood *TRIM13* transcripts was increased in MJD subjects suggesting a peripheral inflammation which may be a reflection of the neuroinflammation found in post-mortem MJD brain.^31,64,65^ *TRIM13* transcript levels inversely correlated with the expanded CAG repeat in the combined group of Azorean and Brazilian MJD patients, and in the subgroup of early-stage Azorean patients *TRIM13* transcript levels inversely correlated with the age of onset. In post-mortem brains, soluble TRIM13 protein levels were increased in MJD patients’ DCN comparing with controls suggesting a potential role of TRIM13 in eliminating misfolded and aggregated proteins in this brain region severely affected by degeneration. Overall, we speculate that the increase of *TRIM13* expression levels in blood and post-mortem DCN samples from MJD subjects could possibly reflect the attempt of cells to promote the clearance of misfolded proteins and aggregates during ER stress and/or to induce apoptosis if the ER stress conditions persist.

*P2RY13* encodes the P2Y purinoceptor 13 protein that belongs to the large G-protein coupled receptors (GPCR) family of membrane proteins.^66^ *P2RY3* is mainly expressed in macrophages cells and in microglial cells.^51^ P2RY13 membrane protein induces the expression and the release of several proinflammatory cytokines.^67^ In this work, blood *P2RY13* transcript levels were increased in Brazilian MJD patients, showing the same dysregulation pattern observed in the Azorean preclinical MJD subjects and early-stage patients. This result is in agreement with those observed in a group of Azorean MJD patients in several stages of the disease up to 30 years after onset.^11^ Interestingly, blood *P2RY13* transcript abundance directly correlated with the age at onset of MJD patients suggesting a link with MJD pathogenesis. Moreover, in this work, insoluble P2RY13 protein levels were increased in patients’ DCN comparing with controls and given that this protein is almost exclusively expressed in microglia in the brain,^51^ this finding suggests a role of this P2Y receptor in neuroinflammation, which is a major player in MJD pathogenesis.^16,31,64,65^ In contrast, soluble P2RY13 protein levels were decreased in patients’ frontal cortex which might indicate a tissue-specific disease mechanism. Overall, the increase of P2RY13 abundance in blood and DCN of MJD subjects may indicate a common inflammatory response in these tissues. Additionally, the fact that the GPCR family member A2aR (adenosine A2a receptor) is associated with MJD-mediated striatal neuropathology in a MJD mouse model^68^ supports the involvement of GPCR family members in the MJD pathogenic process, which have been previously described in other neurodegenerative diseases.^69,70^

Here, the relevance of altered abundance of *DDIT4*, *TRIM13*, and *P2RY13* gene products in MJD subjects is determined by: i) consistent transcriptional behaviors in blood of MJD subjects from two cohorts of different genetic backgrounds, which are markedly distinct from the ones observed in healthy controls; ii) the correlation between transcript levels and patients’ clinical and/or genetic features; and iii) altered DDIT4, TRIM13 and P2RY13 protein levels in MJD brains. Importantly, this study reveals that the abundance of blood *DDIT4* and *TRIM1*3 transcripts differentiated MJD subjects (from preclinical to symptomatic stages) from healthy controls, in particular in the preclinical and early symptomatic stages. Further studies are needed to confirm the potential of these molecules as MJD transcriptional biomarkers, such as testing their behavior with the disease progression in a longitudinal study and assessing how they correlate with other clinical features of the disease (e.g., Scale for the Assessment and Rating of Ataxia (SARA), and Composite Cerebellar Functional Score (CCSF)). In addition, the fact that expression of *DDIT4*, *TRIM13* and *P2RY13* was altered in blood (at the transcript level) and in brain samples (at the protein level) of MJD subjects suggests that these genes/proteins could play a role in the disease pathogenesis given their involvement in various processes known to be dysregulated in MJD, such as inflammation/neuroinflammation. In the future, it would be important to evaluate the abundance of DDIT4, TRIM13 and P2RY13 protein levels in blood samples from MJD subjects as these readouts might provide additional usefulness as biomarkers of this disease.

## Acknowledgements

We are grateful to the MJD subjects and their relatives as well as to the healthy volunteers for participating in this study. The Michigan brain bank at the University of Michigan is partially supported by the NIH/NIA funded Michigan Alzheimer’s Disease Center (P30AG053760 and P30AG072931).

## References

1. Klockgether T, Mariotti C, Paulson HL. Spinocerebellar ataxia. Nat Rev Dis Prim. 2019;5(1):24. doi:10.1038/s41572-019-0074-3

2. Riess O, Rüb U, Pastore A, Bauer P, Schöls L. SCA3: neurological features, pathogenesis and animal models. Cerebellum. 2008;7(2):125–137. doi:10.1007/s12311-008-0013-4

3. Seidel K, Siswanto S, Brunt ERP, Den Dunnen W, Korf HW, Rüb U. Brain pathology of spinocerebellar ataxias. Acta Neuropathol. 2012;124(1):1–21. doi:10.1007/s00401-012-1000-x

4. Maciel P, Costa MC, Ferro A, et al. Improvement in the molecular diagnosis of Machado-Joseph disease. Arch Neurol. 2001;58(11):1821–1827. http://www.ncbi.nlm.nih.gov/pubmed/11708990

5. Wu X, Liao X, Zhan Y, et al. Microstructural Alterations in Asymptomatic and Symptomatic Patients with Spinocerebellar Ataxia Type 3: A Tract-Based Spatial Statistics Study. Front Neurol. 2017;8:714. doi:10.3389/fneur.2017.00714

6. Rezende TJR, de Paiva JLR, Martinez ARM, et al. Structural signature of SCA3: From presymptomatic to late disease stages. Ann Neurol. 2018;84(3):401–408. doi:10.1002/ana.25297

7. Globas C, du Montcel ST, Baliko L, et al. Early symptoms in spinocerebellar ataxia type 1, 2, 3, and 6. Mov Disord. 2008;23(15):2232–2238. doi:10.1002/mds.22288

8. Raposo M, Vasconcelos J, Bettencourt C, Kay T, Coutinho P, Lima M. Nystagmus as an early ocular alteration in Machado-Joseph disease (MJD/SCA3). BMC Neurol. 2014;14:17. doi:10.1186/1471-2377-14-17

9. Costa M do C, Paulson HL. Toward understanding Machado–Joseph disease. Prog Neurobiol. 2012;97(2):239–257. doi:10.1016/j.pneurobio.2011.11.006

10. Evers MM, Toonen LJA, Van Roon-Mom WMC. Ataxin-3 protein and RNA toxicity in spinocerebellar ataxia type 3: Current insights and emerging therapeutic strategies. Mol Neurobiol. 2014;49(3):1513–1531. doi:10.1007/s12035-013-8596-2

11. Raposo M, Bettencourt C, Maciel P, et al. Novel candidate blood-based transcriptional biomarkers of Machado-Joseph disease. Mov Disord. 2015;30(7):968–975. doi:10.1002/mds.26238

12. Kazachkova N, Raposo M, Ramos A, Montiel R, Lima M. Promoter Variant Alters Expression of the Autophagic BECN1 Gene: Implications for Clinical Manifestations of Machado-Joseph Disease. Cerebellum. 2017;16(5-6):957–963. doi:10.1007/s12311-017-0875-4

13. Raposo M, Bettencourt C, Ramos A, et al. Promoter Variation and Expression Levels of Inflammatory Genes IL1A, IL1B, IL6 and TNF in Blood of Spinocerebellar Ataxia Type 3 (SCA3) Patients. NeuroMolecular Med. 2017;19(1):41–45. doi:10.1007/s12017-016-8416-8

14. Raposo M, Hübener-Schmid J, Ferreira AF, et al. Blood transcriptome sequencing identifies biomarkers able to track disease stages in spinocerebellar ataxia type 3. Brain. Published online April 2023. doi:10.1093/brain/awad128

15. Ferreira AF, Raposo M, Shaw ED, et al. Tissue-Specific Vulnerability to Apoptosis in Machado-Joseph Disease. Cells. 2023;12(10). doi:10.3390/cells12101404

16. da Silva Carvalho G, Saute JAM, Haas CB, et al. Cytokines in Machado Joseph Disease/Spinocerebellar Ataxia 3. Cerebellum. 2016;15(4):518–525. doi:10.1007/s12311-015-0719-z

17. de Assis AM, Saute JAM, Longoni A, et al. Peripheral oxidative stress biomarkers in spinocerebellar ataxia type 3/Machado-Joseph disease. Front Neurol. 2017;8(SEP):1–8. doi:10.3389/fneur.2017.00485

18. Raposo M, Ramos A, Santos C, et al. Accumulation of Mitochondrial DNA Common Deletion Since The Preataxic Stage of Machado-Joseph Disease. Mol Neurobiol. 2019;56(1):119–124. doi:10.1007/s12035-018-1069-x

19. Rubinsztein DC, Orr HT. Diminishing return for mechanistic therapeutics with neurodegenerative disease duration?: There may be a point in the course of a neurodegenerative condition where therapeutics targeting disease-causing mechanisms are futile. Bioessays. 2016;38(10):977–980. doi:10.1002/bies.201600048

20. Maas RPPWM, Teerenstra S, Lima M, et al. Differential Temporal Dynamics of Axial and Appendicular Ataxia in SCA3. Mov Disord. 2022;37(9):1850–1860. doi:10.1002/mds.29135

21. Klockgether T, Lüdtke R, Kramer B, Abele M, Bürk K, Schöls L, Riess O, Laccone F, Boesch S, Lopes-Cendes I, Brice A, Inzelberg R, Zilber N DJ. The natural history of degenerative ataxia: a retrospective study in 466 patients. Brain. 1998;121(4):589–600. doi:10.1093/brain/121.4.589

22. Kieling C, Rieder CRM, Silva ACF, et al. A neurological examination score for the assessment of spinocerebellar ataxia 3 (SCA3). Eur J Neurol. 2008;15(4):371–376. doi:10.1111/j.1468-1331.2008.02078.x

23. Raposo M, Ramos A, Bettencourt C, Lima M. Replicating studies of genetic modifiers in spinocerebellar ataxia type 3: Can homogeneous cohorts aid? Brain. 2015;138(12):e398. doi:10.1093/brain/awv206

24. Bettencourt C, Fialho RN, Santos C, et al. Segregation distortion of wild-type alleles at the Machado-Joseph disease locus: a study in normal families from the Azores islands (Portugal). J Hum Genet. 2008;53(4):333–339. doi:10.1007/s10038-008-0261-7

25. Rüb U, Brunt ER, Deller T. New insights into the pathoanatomy of spinocerebellar ataxia type 3 (Machado-Joseph disease). Curr Opin Neurol. 2008;21(2):111–116. doi:10.1097/WCO.0b013e3282f7673d

26. de Rezende TJR, D’Abreu A, Guimaraes RP, et al. Cerebral cortex involvement in Machado-Joseph disease. Eur J Neurol. 2015;22(2):277–283, e23-4. doi:10.1111/ene.12559

27. Arruda WO, Meira AT, Ono SE, et al. Volumetric MRI Changes in Spinocerebellar Ataxia (SCA3 and SCA10) Patients. Cerebellum. 2020;19(4):536–543. doi:10.1007/s12311-020-01137-3

28. Ashraf NS, Duarte-Silva S, Shaw ED, et al. Citalopram Reduces Aggregation of ATXN3 in a YAC Transgenic Mouse Model of Machado-Joseph Disease. Mol Neurobiol. 2019;56(5):3690–3701. doi:10.1007/s12035-018-1331-2

29. Petryszak R, Keays M, Tang YA, et al. Expression Atlas update—an integrated database of gene and protein expression in humans, animals and plants. Nucleic Acids Res. 2015;44(D1):D746–D752. doi:10.1093/nar/gkv1045

30. MacDonald JR, Ziman R, Yuen RKC, Feuk L, Scherer SW. The Database of Genomic Variants: a curated collection of structural variation in the human genome. Nucleic Acids Res. 2014;42(Database issue):D986–92. doi:10.1093/nar/gkt958

31. Evert BO, Vogt IR, Kindermann C, et al. Inflammatory genes are upregulated in expanded ataxin-3-expressing cell lines and spinocerebellar ataxia type 3 brains. J Neurosci. 2001;21(15):5389–5396. doi:10.1523/jneurosci.21-15-05389.2001

32. Hong S, Kim SJ, Ka S, Choi I, Kang S. USP7, a ubiquitin-specific protease, interacts with ataxin-1, the SCA1 gene product. Mol Cell Neurosci. 2002;20(2):298–306. doi:10.1006/mcne.2002.1103

33. Silvestroni A, Faull RLM, Strand AD, Möller T. Distinct neuroinflammatory profile in post-mortem human Huntington’s disease. Neuroreport. 2009;20(12):1098–1103. doi:10.1097/WNR.0b013e32832e34ee

34. Guo L, Giasson BI, Glavis-Bloom A, et al. A cellular system that degrades misfolded proteins and protects against neurodegeneration. Mol Cell. 2014;55(1):15–30. doi:10.1016/j.molcel.2014.04.030

35. He WT, Zheng XM, Zhang YH, et al. Cytoplasmic Ubiquitin-Specific Protease 19 (USP19) Modulates Aggregation of Polyglutamine-Expanded Ataxin-3 and Huntingtin through the HSP90 Chaperone. PLoS One. 2016;11(1):e0147515. doi:10.1371/journal.pone.0147515

36. Zhu G, Harischandra DS, Ghaisas S, et al. TRIM11 Prevents and Reverses Protein Aggregation and Rescues a Mouse Model of Parkinson’s Disease. Cell Rep. 2020;33(9):108418. doi:10.1016/j.celrep.2020.108418

37. Zhang ZY, Harischandra DS, Wang R, et al. TRIM11 protects against tauopathies and is down-regulated in Alzheimer’s disease. Science (80-). 2023;381(6656):eadd6696. doi:10.1126/science.add6696

38. Dietzsch J, Gehlenborg N, Nieselt K. Mayday-a microarray data analysis workbench. Bioinformatics. 2006;22(8):1010–1012. doi:10.1093/bioinformatics/btl070

39. Ferreira AF, Raposo M, Vasconcelos J, Costa M do C, Lima M. Selection of Reference Genes for Normalization of Gene Expression Data in Blood of Machado-Joseph Disease/Spinocerebellar Ataxia Type 3 (MJD/SCA3) Subjects. J Mol Neurosci. 2019;3(69):450–455. doi:10.1007/s12031-019-01374-0

40. Livak KJ, Schmittgen TD. Analysis of Relative Gene Expression Data Using Real-Time Quantitative PCR and the 2−ΔΔCT Method. Methods. 2001;25(4):402–408. doi:10.1006/meth.2001.1262

41. Paulson HL, Das SS, Crino PB, et al. Machado-Joseph disease gene product is a cytoplasmic protein widely expressed in brain. Ann Neurol. 1997;41(4):453–462. doi:10.1002/ana.410410408

42. Youden WJ. Index for rating diagnostic tests. Cancer. 1950;3(1):32–35. doi:10.1002/1097-0142(1950)3:1<32::AID-CNCR2820030106>3.0.CO;2-3

43. Li Q fu, Dong Y, Yang L, et al. Neurofilament light chain is a promising serum biomarker in spinocerebellar ataxia type 3. Mol Neurodegener. 2019;19(39). 10.1186/s13024-019-0338-0

44. Peng Y, Zhang Y, Chen Z, et al. Association of serum neurofilament light and disease severity in patients with spinocerebellar ataxia type 3. Neurology. 2020;95(22):e2977–e2987. doi:10.1212/WNL.0000000000010671

45. Wilke C, Haas E, Reetz K, et al. Neurofilaments in spinocerebellar ataxia type 3: blood biomarkers at the preataxic and ataxic stage in humans and mice. EMBO Mol Med. 2020;12(7):e11803. doi:10.15252/emmm.201911803

46. Prudencio M, Garcia-Moreno H, Jansen-West KR, et al. Toward allele-specific targeting therapy and pharmacodynamic marker for spinocerebellar ataxia type 3. Sci Transl Med. 2020;12(566). doi:10.1126/scitranslmed.abb7086

47. Garcia-Moreno H, Prudencio M, Thomas-Black G, et al. Tau and neurofilament light-chain as fluid biomarkers in spinocerebellar ataxia type 3. Eur J Neurol. 2022;29(8):2439–2452. 10.1111/ene.15373

48. Gonsior K, Kaucher GA, Pelz P, et al. PolyQ-expanded ataxin-3 protein levels in peripheral blood mononuclear cells correlate with clinical parameters in SCA3: a pilot study. J Neurol. 2021;268(4):1304–1315. doi:10.1007/s00415-020-10274-y

49. Hübener-Schmid J, Kuhlbrodt K, Peladan J, et al. Polyglutamine-Expanded Ataxin-3: A Target Engagement Marker for Spinocerebellar Ataxia Type 3 in Peripheral Blood. Mov Disord. 2021;36(11):2675–2681. doi:10.1002/mds.28749

50. Barro C, Chitnis T, Weiner HL. Blood neurofilament light: a critical review of its application to neurologic disease. Ann Clin Transl Neurol. 2020;7(12):2508–2523. doi:10.1002/acn3.51234

51. Karlsson M, Zhang C, Méar L, et al. A single-cell type transcriptomics map of human tissues. Sci Adv. 2021;7(31). doi:10.1126/sciadv.abh2169

52. Corradetti MN, Inoki K, Guan KL. The stress-inducted proteins RTP801 and RTP801L are negative regulators of the mammalian target of rapamycin pathway. J Biol Chem. 2005;280(11):9769–9772. doi:10.1074/jbc.C400557200

53. Sofer A, Lei K, Johannessen CM, Ellisen LW. Regulation of mTOR and cell growth in response to energy stress by REDD1. Mol Cell Biol. 2005;25(14):5834–5845. doi:10.1128/MCB.25.14.5834-5845.2005

54. Malagelada C, Ryu EJ, Biswas SC, Jackson-Lewis V, Greene LA. RTP801 is elevated in Parkinson brain substantia nigral neurons and mediates death in cellular models of Parkinson’s disease by a mechanism involving mammalian target of rapamycin inactivation. J Neurosci. 2006;26(39):9996–10005. doi:10.1523/JNEUROSCI.3292-06.2006

55. Britto FA, Dumas K, Giorgetti-Peraldi S, Ollendorff V, Favier FB. Is REDD1 a metabolic double agent? Lessons from physiology and pathology. Am J Physiol Cell Physiol. 2020;319(5):C807–C824. doi:10.1152/ajpcell.00340.2020

56. Ip WKE, Hoshi N, Shouval DS, Snapper S, Medzhitov R. Anti-inflammatory effect of IL-10 mediated by metabolic reprogramming of macrophages. Science (80-). 2017;356(6337):513–519. doi:10.1126/science.aal3535

57. Pan X, Zhang Z, Liu C, et al. Circulating levels of DDIT4 and mTOR, and contributions of BMI, inflammation and insulin sensitivity in hyperlipidemia. Exp Ther Med. 2022;24(5):666. doi:10.3892/etm.2022.11602

58. Huang B, Baek SH. Trim13 Potentiates Toll-Like Receptor 2-Mediated Nuclear Factor kappaB Activation via K29-Linked Polyubiquitination of Tumor Necrosis Factor Receptor-Associated Factor 6. Mol Pharmacol. 2017;91(4):307–316. doi:10.1124/mol.116.106716

59. Mikael Lerner, Martin Corcoran, Diana Cepeda, Michael L. Nielsen, Roman Zubarev, Fredrik Pontén, Mathias Uhlén, Sophia Hober, Dan Grandér OS. The RBCC Gene RFP2 (Leu5) Encodes a Novel Transmembrane E3 Ubiquitin Ligase Involved in ERAD. Mol Biol Cell. 2007;18:1670–1682. doi:10.1091/mbc.E06-03-0248

60. Tomar D, Singh R, Singh AK, Pandya CD, Singh R. TRIM13 regulates ER stress induced autophagy and clonogenic ability of the cells. Biochim Biophys Acta - Mol Cell Res. 2012;1823(2):316–326. doi:10.1016/j.bbamcr.2011.11.015

61. Ji CH, Kim HY, Heo AJ, et al. The N-Degron Pathway Mediates ER-phagy. Mol Cell. 2019;75(5):1058–1072.e9. doi:10.1016/j.molcel.2019.06.028

62. Tomar D, Prajapati P, Sripada L, et al. TRIM13 regulates caspase-8 ubiquitination, translocation to autophagosomes and activation during ER stress induced cell death. Biochim Biophys Acta. 2013;1833(12):3134–3144. doi:10.1016/j.bbamcr.2013.08.021

63. Zhang L, Afolabi LO, Wan X, Li Y, Chen L. Emerging Roles of Tripartite Motif-Containing Family Proteins (TRIMs) in Eliminating Misfolded Proteins. Front Cell Dev Biol. 2020;8:802. https://www.frontiersin.org/article/10.3389/fcell.2020.00802

64. Evert BO, Schelhaas J, Fleischer H, et al. Neuronal intranuclear inclusions, dysregulation of cytokine expression and cell death in spinocerebellar ataxia type 3. Clin Neuropathol. 2006;25(6):272–281.

65. Duarte Lobo D, Nobre RJ, Oliveira Miranda C, et al. The blood-brain barrier is disrupted in Machado-Joseph disease/spinocerebellar ataxia type 3: evidence from transgenic mice and human post-mortem samples. Acta Neuropathol Commun. 2020;8(1):152. doi:10.1186/s40478-020-00955-0

66. Communi D, Gonzalez NS, Detheux M, et al. Identification of a novel human ADP receptor coupled to G(i). J Biol Chem. 2001;276(44):41479–41485. doi:10.1074/jbc.M105912200

67. Liu PW, Yue MX, Zhou R, et al. P2Y(12) and P2Y(13) receptors involved in ADPβs induced the release of IL-1β, IL-6 and TNF-α from cultured dorsal horn microglia. J Pain Res. 2017;10:1755–1767. doi:10.2147/JPR.S137131

68. Gonçalves N, Simões AT, Cunha RA, De Almeida LP. Caffeine and adenosine A2A receptor inactivation decrease striatal neuropathology in a lentiviral-based model of Machado-Joseph disease. Ann Neurol. 2013;73(5):655–666. doi:10.1002/ana.23866

69. Huang Y, Todd N, Thathiah A. The role of GPCRs in neurodegenerative diseases: avenues for therapeutic intervention. Curr Opin Pharmacol. 2017;32:96–110. doi:10.1016/j.coph.2017.02.001

70. Azam S, Haque ME, Jakaria M, Jo SH, Kim IS, Choi DK. G-Protein-Coupled Receptors in CNS: A Potential Therapeutic Target for Intervention in Neurodegenerative Disorders and Associated Cognitive Deficits. Cells. 2020;9(2):506. doi:10.3390/cells9020506

71. Krämer A, Green J, Pollard J, Tugendreich S. Causal analysis approaches in ingenuity pathway analysis. Bioinformatics. 2014;30(4):523–530. doi:10.1093/bioinformatics/btt703

