## Supplementary Tables 1-7 and Figure 1 for "Blood *DDIT4* and *TRIM13* transcript levels mark the early stages of Machado-Joseph disease"

### Supplementary data

**Supplementary Table 1 Characterization of the MJD subjects and control individuals used in this study**

| | Microarray analysis/ technical validation | Validation in independent cohorts | | Early disease stage | Post-mortem human brain |
|---|---|---|---|---|---|
| | **Group #1 (Azores)** | **Group #2 (Azores)** | **Group #3 (Brazil)** | **Group #4 (Azores)** | |
| **Preclinical subjects** | | | | | |
| N (Female, Male) | 9 (7, 2) | 10 (6, 4) | NA | 19 (12, 7)[f] | NA |
| Age[a], years | 27.9 ± 6.9 [22; 43] | 32.5 ± 7.4 [21; 44] | NA | 30.3 ± 7.3 [21; 44] | NA |
| $CAG_n$ allele 1[b] | 19.6 ± 4.6 [14; 28] | 19.1 ± 3.7 [14; 23] | NA | 20.0 ± 4.1 [14; 28] | NA |
| $CAG_n$ allele 2[c] | 68.4 ± 2.9 [65; 74] | 67.5 ± 3.3 [62; 75] | NA | 68.1 ± 3.0 [62;75] | NA |
| Years to onset[d] | -12.4 ± 7.9 [-27; -4] | -9.5 ± 10.4 [-25; +5] | NA | 10.4 ± 9.3 [-26; +5] | NA |
| **Patients** | | | | | |
| N (Female, Male) | 11 (4, 7) | 28 (15, 13) | 20 (8, 12) | 36 (17, 19) | 5 (4, 1) |
| Age[a], years | 54.6 ± 10.7 [37; 72] | 47.6 ± 14.3 [25; 85] | 42.7 ± 8.1 [30; 58] | 39.8 ± 10.9 [17; 63] | NA |
| $CAG_n$ allele 1[b] | 21.2 ± 5.1 [14; 27] | 22.0 ± 4.6 [14; 29] | 21.3 ± 4.7 [14; 29] | 22.0 ± 5.0 [14; 29] | 19.0 ± 4.1 [12; 22] |
| $CAG_n$ allele 2[c] | 69.4 ± 2.8 [65; 74] | 70.0 ± 3.8 [60; 76] | 74.7 ± 2.8 [70; 80] | 71.3 ± 3.4 [65;79] | 70.2 ± 3.0 [66; 73] |
| Age at onset, years | 42.5 ± 12.1 [25; 60] | 37.7 ± 11.3 [22; 70] | 35.8 ± 7.0 [25; 47] | 37.2 ± 11.0 [16; 60] | 45.0 ± 8.5 [39; 51][h] |
| Disease duration, years | 12.2 ± 10.5 [2; 27] | 9.9 ± 7.4 [0; 25] | 6.9 ± 2.3 [1; 11] | 2.7 ± 1.5 [0; 5] | 26.5 ± 9.2 [20; 33][h] |
| NESSCA | 11.5 ± 5.0 [6; 22] | 12.5 ± 4.2 [4; 24] | 14.9 ± 5.0 [8; 27] | 8.7 ± 2.3 [4; 12][g] | NA |
| Age at death, years | NA | NA | NA | NA | 63.0 ± 16.0 [48; 84] |
| PMI[e], hours | NA | NA | NA | NA | 26.6 ± 17.2 [4; 48] |

**Supplementary Table 1 (cont.)**

| | Microarray analysis/ technical validation | Validation in independent cohorts | | Early disease stage | Post-mortem human brain |
|---|---|---|---|---|---|
| | Group #1 (Azores) | Group #2 (Azores) | Group #3 (Brazil) | Group #4 (Azores) | |
| **Controls** | | | | | |
| N (Female, Male) | 8 (5, 3) | 20 (10, 10) | 20 (11, 9) | 55 (29, 26) | 9 (5, 4) |
| Age[a], years | 36.3 ± 11.7 [20; 53] | 42.7 ± 16.0 [22; 77] | 40.3 ± 12.7 [27; 69] | matched paired | NA |
| $CAG_n$ allele 1[b] | NA | NA | NA | 19.1 ± 4.0 [15; 27] | 17.0 ± 5.3 [11; 25] |
| $CAG_n$ allele 2[c] | NA | NA | NA | 23.6 ± 3.9 [15; 32] | 22.6 ± 5.6 [12; 30] |
| Age at death, years | NA | NA | NA | NA | 69.6 ± 12.2 [48; 83] |
| PMI[e], hours | NA | NA | NA | NA | 15.4 ± 8.0 [4; 24] |

Quantitative variables are displayed as mean ± standard deviation [minimum; maximum]; [a]Age at blood collection, [b]Number of CAG repeats in the normal allele of MJD subjects/number of CAG repeats in normal allele 1 of controls; [c]Number of CAG repeats in expanded allele of MJD subjects/number of CAG repeats in normal allele 2 of controls; [d]Years to onset: negative values indicate the numbers of years missing to the estimated onset and positive values indicate the number of years that have elapsed the estimated onset; [e]Post-mortem interval; [p]reclinical subjects comprised 18 from group #1-2, and one additional individual; [g]Information available for only 14 patients; [h]Information available for two patients; NA, not applicable/not available

**Supplementary Table 2 Characterization of post-mortem human brain samples from MJD patients and controls individuals, and RNA integrity number of each brain samples used in this study**

| Health condition | Sex | ID | Age at death (years) | PMI[a] (hours) | Cause of death | Age at onset (years) | CAG repeats | RNA integrity number | | |
|---|---|---|---|---|---|---|---|---|---|---|
| | | | | | | | Allele 1 \| Allele 2 | DCN[b] | Pons | Frontal cortex |
| Controls | Female | 1 | 48 | 5 | Polycythemia vera, mesenteric thyrombosis and ischemic bowel resection | NA | 12 \| 19 | 3.7 | 3.2 | 3.1 |
| | | 2 | 76 | 14 | Cardiac failure | NA | 21 \| 25 | 2.9 | 5.3 | 3.4 |
| | | 3 | 80 | 19 | Congestive heart failure and atrial fibrillation | NA | 12 \| 18 | 5.2 | 6 | NA |
| | | 4 | 83 | NA | Renal cell carcinoma | NA | 18 \| 21 | 4.8 | 4.6 | 5.6 |
| | | 5 | 83 | 21 | Cardiac arrest, urinary tract infection and sepsis | NA | 11 \| 12 | 6.9 | 5.7 | NA |
| | Male | 6 | 59 | 12 | Sudden cardiac arrest, ventricular fibrillation and post-shock electromechanical dissociation | NA | 12 \| 25 | 7.7 | NA | 6.5 |
| | | 7 | 61 | 24 | Cardiac failure, cardiogenic shock and post-shock electromechanical dissociation | NA | 21 \| 25 | 7.5 | NA | 6.9 |
| | | 8 | 65 | 24 | Acute respiratory distress syndrome and sepsis | NA | 25 \| 28 | NA | NA | NA |
| | | 9 | 71 | 4 | Cardiac failure | NA | 21 \| 30 | 7 | 7.4 | 8.3 |
| MJD patients | Female | 10 | 48 | 22 | NA | NA | 22 \| 73 | 5.3 | 4.1 | 6.2 |
| | | 11 | 59 | 4 | NA | 39 | 21 \| 70 | 7.9 | 6.2 | 8.4 |
| | | 12 | 75 | 39 | NA | NA | 19 \| 69 | NA | NA | 4 |
| | | 13 | 84 | 20 | NA | 51 | 21 \| 66 | 6.5 | NA | 4.7 |
| | Male | 14 | 49 | 48 | NA | NA | 12 \| 73 | NA | 3.5 | 3.9 |

[a]Post-mortem interval; [b]Dentate cerebellar nucleus; NA, not applicable/not available

**Supplementary Table 3 Microarray data of the genes selected for subsequent qPCR validation**

| Gene Symbol[a] | Gene Name[a] | Preclinical subjects vs. Controls | | Patients vs. Controls | | Patients vs. Preclinical subjects | |
|---|---|---|---|---|---|---|---|
| | | Log ratio | *q* value | Log ratio | *q* value | Log ratio | *q* value |
| **Enzimes[b]** | | | | | | | |
| *CLC* | Charcot-Leyden crystal galectin | -0.560 | 0.596 | 0.788 | **0.000** | 1.348 | **0.000** |
| *FKBP14* | FKBP prolyl isomerase 14 | 0.684 | **0.000** | 0.868 | **0.000** | 0.183 | 0.305 |
| *HSD17B7* | hydroxysteroid 17-beta dehydrogenase 7 | 0.634 | **0.000** | 0.753 | **0.000** | 0.119 | 1.000 |
| *LYZ* | lysozyme | -0.367 | 0.607 | 0.009 | 0.498 | 0.376 | **0.000** |
| *MAGT1* | magnesium transporter 1 | 0.822 | **0.000** | 0.795 | **0.000** | -0.028 | 0.880 |
| *N4BP2* | NEDD4 binding protein 2 | 0.341 | 1.000 | 0.462 | **0.015** | 0.121 | 1.000 |
| *PGGHG* | protein-glucosylgalactosylhydroxylysine glucosidase | 1.050 | **0.000** | 0.716 | **0.000** | -0.335 | 1.000 |
| *TRIM13* | tripartite motif containing 13 | 0.350 | 1.000 | 0.502 | **0.015** | 0.152 | 1.000 |
| **Kinases** | | | | | | | |
| *EIF2AK4* | eukaryotic translation initiation factor 2 alpha kinase 4 | 0.269 | 1.000 | 0.483 | **0.015** | 0.215 | 1.000 |
| *PIP4K2B* | phosphatidylinositol-5-phosphate 4-kinase type 2 beta | 0.427 | 0.223 | 0.651 | **0.000** | 0.223 | 0.723 |
| *SGK1* | serum/glucocorticoid regulated kinase 1 | 0.344 | 1.000 | 0.566 | **0.000** | 0.222 | 1.000 |
| **Peptidases** | | | | | | | |
| *GZMH* | granzyme H | -0.806 | **0.000** | -0.560 | 0.124 | 0.246 | 0.521 |
| *MMP9* | matrix metallopeptidase 9 | 0.310 | 1.000 | -0.121 | 0.398 | -0.431 | **0.000** |
| *USP49* | ubiquitin specific peptidase 49 | 0.450 | 0.596 | 0.657 | **0.000** | 0.207 | 0.723 |
| **Transcription regulators** | | | | | | | |
| *BLZF1* | basic leucine zipper nuclear factor 1 | 0.547 | 0.221 | 0.719 | **0.000** | 0.172 | 0.590 |
| *CREB1* | cAMP responsive element binding protein 1 | 0.509 | 0.478 | 0.595 | **0.000** | 0.086 | 1.000 |
| **Transmembrane receptor** | | | | | | | |
| *FCER1A* | Fc fragment of IgE receptor Ia | 0.155 | 1.000 | 0.797 | **0.000** | 0.642 | **0.000** |

**Supplementary Table 3 (cont.)**

| Gene Symbol[a] | Gene Name[a] | Preclinical subjects vs. Controls | | Patients vs. Controls | | Patients vs. Preclinical subjects | |
|---|---|---|---|---|---|---|---|
| | | Log ratio | *q* value | Log ratio | *q* value | Log ratio | *q* value |
| **Others[b]** | | | | | | | |
| *CCDC125* | coiled-coil domain containing 125 | 0.436 | 0.358 | 0.676 | **0.000** | 0.241 | 0.593 |
| *DDIT4* | DNA damage inducible transcript 4 | -0.595 | 0.221 | -0.623 | **0.000** | -0.028 | 1.000 |
| *LGALS2* | galectin 2 | -0.747 | **0.000** | -0.373 | 0.342 | 0.374 | 0.142 |
| *PI3* | peptidase inhibitor 3 | -0.633 | **0.000** | -0.428 | **0.000** | 0.205 | 0.903 |
| *S100P* | S100 calcium binding protein P | 0.586 | 0.221 | 0.577 | **0.000** | -0.009 | 0.802 |
| *SHROOM4* | shroom family member 4 | 0.658 | **0.000** | 0.686 | **0.000** | 0.028 | 1.000 |
| *SVBP* | small vasohibin binding protein | 0.326 | 0.221 | -0.164 | 1.000 | -0.490 | **0.000** |

[a]Official gene symbol and full name provided by HUGO Gene Nomenclature Committee; [b]Gene ontology annotations were performed in Ingenuity Pathway Analysis[71]; Down- and upregulated genes are indicated in green and red colors, respectively; Bold values denote statistical significance at a Benjamini-Hochberg false discovery rate (FDR) adjusted *p*-value ($q<0.05$).

**Supplementary Table 4 Genes selected from the microarray analysis for technical and biological validation by qPCR**

| | | | | | | | | | | | | | | | |
|---|---|---|---|---|---|---|---|---|---|---|---|---|---|---|---|
| | | | | | | | | **Validation in independent cohorts** | | | | | | | |
| | | **Technical Validation Group #1 (Azores)[c]** | | | | | | **Group #2 (Azores)[c]** | | | | | | **Group #3 (Brazil)[d]** | |
| | | **PC subjects vs. Controls** | | **Patients vs. Controls** | | **Patients vs. PC subjects** | | **PC subjects vs. Controls** | | **Patients vs. Controls** | | **Patients vs. PC subjects** | | **Patients vs. Controls** | |
| **Gene Symbol[a]** | **TaqMan Assay ID** | **Mean diff.** | ***p* value** | **Mean diff.** | ***p* value** | **Mean diff.** | ***p* value** | **Mean diff.** | ***p* value** | **Mean diff.** | ***p* value** | **Mean dif.** | ***p* value** | **Mean diff.** | ***p* value** |
| **Enzimes[b]** | | | | | | | | | | | | | | | |
| *CLC* | Hs01055743_m1 | 0.039 | 1.000 | 0.194 | 0.409 | 0.155 | 0.940 | | | | | | | | |
| *FKBP14* | Hs00215735_m1 | 0.012 | 0.294 | 0.010 | 0.409 | -0.002 | 1.000 | | | | | | | | |
| *HSD17B7* | Hs00367686_m1 | 0.012 | **0.000** | 0.003 | 0.919 | -0.008 | 0.065 | 0.048 | **0.008** | -0.012 | 0.109 | -0.060 | **0.001** | | |
| *LYZ* | Hs00426232_m1 | -2.261 | 1.000 | 0.863 | 1.000 | 3.123 | 1.000 | | | | | | | | |
| *MAGT1* | Hs00259564_m1 | 0.023 | 0.849 | -0.014 | 1.000 | -0.037 | 0.618 | | | | | | | | |
| *N4BP2* | Hs00905983_m1 | 0.009 | 0.085 | -0.004 | 1.000 | -0.013 | 0.060 | | | | | | | | |
| *PGGHG* | HS00228253_m1 | 0.287 | **0.000** | 0.220 | **0.029** | -0.066 | 1.000 | 0.172 | 0.564 | -0.053 | 1.000 | -0.224 | 0.247 | | |
| *TRIM13* | Hs00328634_s1 | 0.024 | 0.683 | 0.024 | 0.930 | -0.001 | 1.000 | 0.156 | **0.000** | 0.084 | **0.000** | -0.073 | 0.281 | 0.044 | **0.004** |
| **Kinases** | | | | | | | | | | | | | | | |
| *EIF2AK4* | Hs01010957_m1 | 0.019 | 0.407 | -0.020 | 0.540 | -0.039 | 0.077 | | | | | | | | |
| *PIP4K2B* | Hs01594707_m1 | 0.018 | 0.536 | -0.008 | 1.000 | -0.026 | 0.476 | | | | | | | | |
| *SGK1* | Hs00985033_g1 | 0.084 | 0.592 | 0.144 | 0.162 | 0.061 | 1.000 | | | | | | | | |
| **Peptidases** | | | | | | | | | | | | | | | |
| *GZMH* | Hs00277212_m1 | -0.198 | 0.758 | -0.440 | 0.082 | -0.242 | 0.911 | | | | | | | | |
| *MMP9* | Hs00234579_m1 | 0.276 | 1.000 | -0.396 | 0.243 | -0.672 | 0.316 | 0.463 | 0.115 | 0.182 | 0.209 | -0.282 | 0.703 | | |
| *USP49* | Hs00988691_m1 | 0.005 | 0.380 | 0.006 | 0.251 | 0.001 | 1.000 | 0.023 | 0.885 | -0.040 | **0.000** | -0.063 | **0.005** | 0.011 | 0.064 |
| **Transcription regulators** | | | | | | | | | | | | | | | |
| *BLZF1* | Hs00388707_m1 | 0.020 | **0.008** | 0.007 | 1.000 | -0.012 | 0.477 | 0.057 | **0.002** | -0.004 | 1.000 | -0.061 | **0.001** | | |
| *CREB1* | Hs00231713_m1 | 0.174 | **0.001** | 0.012 | 1.000 | -0.162 | **0.043** | 0.243 | **0.000** | 0.069 | 0.120 | -0.174 | **0.026** | | |
| **Transmembrane receptor** | | | | | | | | | | | | | | | |
| *FCER1A* | Hs00758600_m1 | 0.025 | 0.176 | 0.054 | **0.001** | 0.029 | 0.306 | 0.026 | 1.000 | 0.006 | 1.000 | -0.020 | 1.000 | | |

**Supplementary Table 4 (cont.)**

| | | | | | | | | Validation in independent cohorts | | | | | | | |
|---|---|---|---|---|---|---|---|---|---|---|---|---|---|---|---|
| | | Technical Validation Group #1 (Azores)[c] | | | | | | Group #2 (Azores)[c] | | | | | | Group #3 (Brazil)[d] | |
| | | PC subjects vs. Controls | | Patients vs. Controls | | Patients vs. PC subjects | | PC subjects vs. Controls | | Patients vs. Controls | | Patients vs. PC subjects | | Patients vs. Controls | |
| **Gene Symbol[a]** | **TaqMan Assay ID** | **Mean diff.** | ***p* value** | **Mean diff.** | ***p* value** | **Mean diff.** | ***p* value** | **Mean diff.** | ***p* value** | **Mean diff.** | ***p* value** | **Mean dif.** | ***p* value** | **Mean diff.** | ***p* value** |
| **Others[b]** | | | | | | | | | | | | | | | |
| *CCDC125* | Hs00417596_m1 | 0.063 | **0.028** | 0.028 | 0.355 | -0.035 | 0.932 | 0.061 | **0.009** | 0.002 | 1.000 | -0.060 | **0.016** | | |
| *DDIT4* | Hs01111686_g1 | -0.148 | 0.059 | -0.219 | **0.008** | -0.071 | 1.000 | -0.056 | 1.000 | -0.139 | **0.004** | -0.083 | 0.357 | -0.071 | **0.002** |
| *LGALS2* | Hs00197810_m1 | -0.281 | 0.091 | -0.306 | 0.099 | -0.025 | 1.000 | | | | | | | | |
| *PI3* | Hs00160066_m1 | -0.045 | 1.000 | -0.152 | 0.345 | -0.107 | 1.000 | | | | | | | | |
| *S100P* | Hs00195584_m1 | 0.605 | 0.080 | 0.011 | 1.000 | -0.594 | 0.330 | 0.475 | 0.051 | 0.320 | **0.008** | -0.155 | 1.000 | -0.094 | 0.256 |
| *SHROOM4* | Hs00292474_m1 | 0.002 | 1.000 | -0.003 | 0.598 | -0.006 | 0.281 | | | | | | | | |
| *SVBP* | Hs00380672_m1 | 0.208 | **0.013** | -0.051 | 0.422 | -0.260 | **0.004** | 0.145 | **0.007** | 0.081 | **0.007** | -0.064 | 0.655 | -0.344 | **0.004** |

[a]Official gene symbol and full name provided by HUGO Gene Nomenclature Committee; [b] Gene ontology annotations were performed in Ingenuity Pathway Analysis[71]; [c]The analysis were performed using a Generalized Linear Model with Bonferroni correction for age at blood collection, and the results are indicated as the estimated mean differences; [d]The analysis were performed using a Mann-Whitney Test. The results are indicated as the mean differences between patients and controls; Down- and upregulated genes are indicated in green and red colors, respectively; Bold values denote statistical significance at a $p<0.05$; PC = preclinical subjects

**Supplementary Table 5 Correlations of transcript levels of *DDIT4*, *TRIM13* and *P2RY13* with genetic and clinical features of MJD patients**

| | | Expanded CAG repeat (CAG-E) | Age at onset[a] | Disease duration (DD) | | Total NESSCA |
|---|---|---|---|---|---|---|
| | | | | no adj. | CAG-E adj. | CAG-E + DD adj. |
| ***DDIT4*** | rho | -0.290 | -0.236 | NA | -0.014 | -0.127 |
| | *p*-value | **0.027** | 0.077 | NA | 0.917 | 0.356 |
| ***TRIM13*** | rho | -0.330 | -0.176 | NA | 0.164 | 0.190 |
| | *p*-value | **0.015** | 0.207 | NA | 0.242 | 0.177 |
| ***P2RY13*[b]** | rho | 0.000 | 0.366 | 0.078 | NA | -0.087 |
| | *p*-value | 0.997 | **0.004** | 0.545 | NA | 0.724 |

[a]Correlation of transcript levels with the age at onset was adjusted for the expanded CAG repeat; [b]Patients with NESSCA score available (n=22); NA, not applicable; statistical significance was set at a $p<0.05$ (bold)

**Supplementary Table 6 Correlations between transcript levels and genetic and clinical features of MJD preclinical subjects and early patients**

| | | | Expanded CAG repeat | Years to onset | Age at onset[a] | Disease duration |
|---|---|---|---|---|---|---|
| Preclinical subjects | ***DDIT4*** | rho | -0.114 | 0.232 | NA | NA |
| | | *p*-value | 0.642 | 0.340 | NA | NA |
| | ***TRIM13*** | rho | -0.437 | 0.160 | NA | NA |
| | | *p*-value | 0.061 | 0.514 | NA | NA |
| | ***P2RY13*** | rho | -0.265 | 0.002 | NA | NA |
| | | *p*-value | 0.274 | 0.994 | NA | NA |
| Early patients | ***DDIT4*** | rho | -0.088 | NA | -0.090 | -0.007 |
| | | *p*-value | 0.610 | NA | 0.607 | 0.967 |
| | ***TRIM13*** | rho | -0.274 | NA | -0.472 | 0.261 |
| | | *p*-value | 0.105 | NA | **0.004** | 0.124 |
| | ***P2RY13*** | rho | -0.069 | NA | -0.217 | -0.030 |
| | | *p*-value | 0.700 | NA | 0.224 | 0.864 |

[a]Correlation of transcript levels with the age at onset was adjusted for the expanded CAG repeat; NA, not applicable; statistical significance was set at a $p<0.05$ (bold)

**Supplementary Table 7 Discrimination ability of transcript levels to distinguish MJD subjects (preclinical subjects or patients) from the respective control individuals**

| | AUC (95% CI)[a] | *p*-value | Optimal cut-off[b] | Sensitivity | Specificity |
|---|---|---|---|---|---|
| **Early stages disease (Group #4)** | | | | | |
| **Preclinical subjects vs. Controls** | | | | | |
| *DDIT4* | 0.94 (0.86-1.0) | < 0.001 | < 3.8 | 100 | 78 |
| *TRIM13* | 0.99 (0.97-1.0) | < 0.001 | > 0.48 | 95 | 95 |
| *P2RY13* | 0.78 (0.63-0.93) | 0.004 | > 2.80 | 84 | 68 |
| *DDIT4* and *TRIM13* | 1.0 (1.0-1.0) | < 0.001 | > 0.50 | 100 | 100 |
| **Early patients vs. Controls** | | | | | |
| *DDIT4* | 0.93 (0.87-0.98) | < 0.001 | < 3.70 | 97 | 75 |
| | | | < 4.10 | 100 | 72 |
| *TRIM13* | 0.92 (0.85-0.98) | < 0.001 | > 0.57 | 89 | 80 |
| *P2RY13* | 0.68 (0.55-0.81) | 0.01 | > 2.8 | 76 | 56 |
| *DDIT4* and *TRIM13* | 0.96 (0.92-1.0) | < 0.001 | > 0.45 | 94 | 89 |
| *DDIT4*, *TRIM13* and *P2RY13* | 0.96 (0.91-1.0) | < 0.001 | > 0.47 | 94 | 88 |
| **Patients ≥5 years of disease duration** | | | | | |
| **Patients vs. Controls** | | | | | |
| *DDIT4* | 0.67 (0.56-0.78) | 0.007 | < 0.24 | 90 | 46 |
| *TRIM13* | 0.85 (0.77-0.93) | <0.001 | > 0.097 | 78 | 87 |
| *P2RY13* | 0.69 (0.59-0.80) | <0.001 | > 0.54 | 66 | 71 |
| *DDIT4* and *TRIM13* | 0.90 (0.83-0.96) | <0.001 | < 0.57 | 84 | 83 |
| | | | < 0.60 | 86 | 81 |

[a]Area under the curve and 95% Confidence Interval (CI); [b]Optimal cut-off was defined by the highest value of the Youden Index (sensitivity + specificity − 1) calculated based on the receiver operating characteristic (ROC) curve; statistical significance was set at $p<0.05$

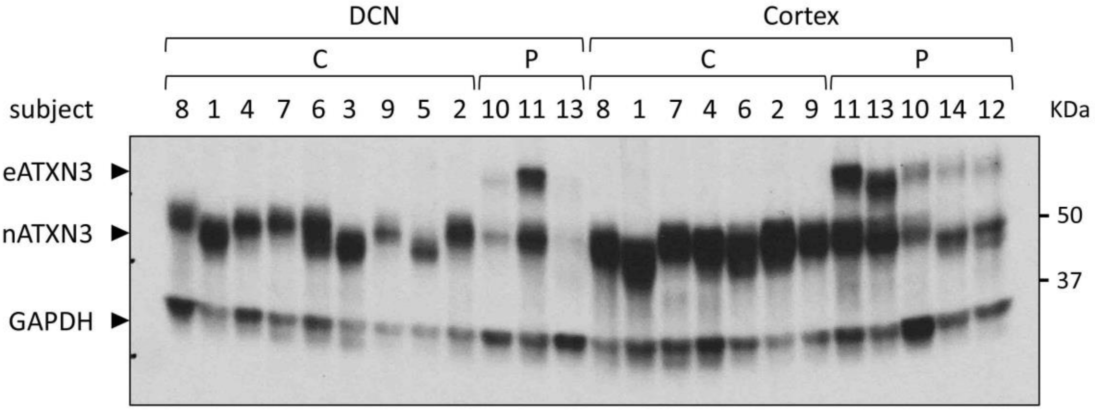


**Supplementary Figure 1** Detection of the native human ATXN3 and the expanded human ATXN3 in post-mortem human samples. Western blot using the anti-MJD antibody[41] to detect the native human ATXN3 (nATXN3) and the expanded human ATXN3 (eATXN3) in insoluble protein fractions of post-mortem human samples from dentate cerebellar nucleus (DCN) and frontal cortex (Cortex) of Machado-Joseph disease patients (P) and control subjects (C). GAPDH was used as a protein loading control.
